# Statistical learning of incidental perceptual regularities induces sensory conditioned cortical responses

**DOI:** 10.1101/2024.02.28.582496

**Authors:** Antonino Greco, Marco D’Alessandro, Giuseppe Gallitto, Clara Rastelli, Christoph Braun, Andrea Caria

**Author notes:** Corresponding authors **Corresponding authors** Antonino Greco, Department of Neural Dynamics and Magnetoencephalography, Hertie Institute for Clinical Brain Research, Otfried-Müller-Straße 47, 72076 Tübingen, Germany, Andrea Caria, Dipartimento di Psicologia e Scienze Cognitive, Corso Bettini, 31, Rovereto, Trento 38068, Italy.

## Abstract

Statistical learning of sensory patterns can lead to predictive neural processes enhancing stimulus perception and enabling fast deviancy detection. Predictive processes have been extensively demonstrated when environmental statistical regularities are relevant to task execution. Preliminary evidence indicates that statistical learning can even occur independently of task relevance and top-down attention, although the temporal profile and neural mechanisms underlying sensory predictions and error signals induced by statistical learning of incidental sensory regularities remain unclear. In our study, we adopted an implicit sensory conditioning paradigm that elicited the generation of specific perceptual priors in relation to task-irrelevant audio-visual associations, while recording Electroencephalography (EEG). Our results showed that learning of non-relevant but statistically interrelated neutral audio-visual stimuli resulted in early neural responses to predictive auditory stimuli conveying anticipatory signals of expected visual stimulus presence or absence, and in specific modulation of cortical responses to probabilistic visual stimulus presentation or omission. Pattern similarity analysis indicated that predictive auditory stimuli tended to resemble the response to expected visual stimulus presence or absence. Remarkably, Hierarchical Gaussian filter modeling estimating dynamic changes of prediction error signals in relation to differential probabilistic occurrences of audio-visual stimuli further demonstrated instantiation of predictive neural signals by showing distinct neural processing of prediction error in relation to violation of expected visual stimulus presence or absence. Overall, our findings indicated that statistical learning of non-salient and task-irrelevant perceptual regularities can induce the generation of neural priors at the time of predictive stimulus presentation, possibly conveying sensory-specific information of the predicted consecutive stimulus.

## 1. INTRODUCTION

Human sensory perception is posited to depend upon rapid encoding of probabilistic occurrence of perceptual information and consequent timely generation of specific sensory priors ^1-6^. A number of studies indicated that statistical learning of sensory patterns leads to predictive neural processes enhancing visual perception and enabling fast deviancy detection ^7-13^. Predictive models of perception postulate that high-level generative neural signals provide anticipatory neural representations of sensory signals reducing perceptual surprisal and minimize computational effort ^3,14-17^. Accordingly, rapid learning of relevant perceptual regularities has been shown to induce attenuation of neural population reaction to predictable sensory stimuli and enhancement of response to unpredictable information ^8,18-21^. The neural suppression effect is proposed to arise either from dampening of stimulus response as result of global surprise signals decrease ^22,23^ or from sharpening of cortical response to specific sensory input and inherent reduction of prediction errors ^9,24^. In line with this last assumption, prior expectations can facilitate sensory processing by increasing fine-tuning of early sensory cortex ^9,19,24,25^. Moreover, encoding of predictable sensory patterns was also associated with anticipatory perceptual processes such as pre-stimulus-specific baseline shifts ^26,27^, and pre-activation of primary sensory cortex, immediately before stimulus presentation, resulting in timely cortical instantiation of the expected stimulus representational content ^28-30^.

Learned perceptual statistical regularities and resulting predictive neural processes are significantly influenced by task-relevance and attention ^28,31-34^. However, statistical learning in the absence of explicit top-down attention can indeed occur and also leads to attentional suppression effect ^35^. In addition, attenuation of cortical fMRI response to predicted stimuli was also observed in case of task-irrelevant sensory input ^8,19,36^.

Stimulus relevance and probability appear to have dissociable effects on visual processing ^13,34^. For instance, stimulus relevance can enhance precision of stimulus processing by suppressing internal noise, whereas sensory signal probability would bias stimulus detection by increasing the baseline activity of signal-selective units during early visual processing ^13^. Notably, learned probability of sensory stimulus occurrence can significantly and differentially impact both late and early response stages ^13,28^.

While the predictive processes associated with responses of predictable information, such as cortical response attenuation, are well documented, the neural mechanisms underlying early processing of predictive and predicted sensory information, in particular when sensory signals are task-irrelevant, still remain unclear. Learning of non-salient sensory regularities can be shaped through associative mechanisms leading to sensory conditioned stimuli capable of inducing anticipatory responses similar to unconditioned responses ^37-40^. Previous sensory pre-conditioning studies typically included an initial sensory conditioning phase followed by Pavlovian conditioning, raising the question whether it was the consecutive conditioning phase involving potent biological stimuli that significantly influenced learned associations between neutral sensory stimuli. On the other hand, functional neuroimaging studies suggested that learning of incidental audio-visual regularities might induce modifications of neural responses to paired sensory inputs independently of stimulus salience ^19,25^.

However, the temporal profile and neural mechanisms underlying sensory predictions and error signals of non-salient and task-irrelevant but statistically-related sensory stimuli remain unclear. In our investigation, on the basis of predictive processing principles we hypothesized that associative learning of non-relevant but probabilistically interrelated neutral sensory stimuli resulted in early neural responses to predictive sensory stimuli associated with anticipatory signals informative of the subsequent sensory input, as well as a specific response modulation to the predicted stimulus. To this aim, we collected electroencephalographic (EEG) data during a novel implicit sensory conditioning paradigm that throughout its unfolding was expected to induce increasingly specific perceptual priors relative to probabilistic but task-irrelevant audio-visual associations. The experimental protocol entailed participants being exposed to incidental probabilistic associations of non-salient audio-visual patterns while engaged in a main stimulus detection task (Figure 1). We specifically expected that implicit learning of task-irrelevant audio-visual associations resulted in anticipatory brain activity associated with auditory stimuli (audio period) conveying information of the consecutive visual stimulus presentation (post-audio period), as possibly evidenced by increased similarity between the pattern of neural response of the predictive auditory stimulus with that of the predicted visual stimulus. We also conjectured that such anticipatory forward mechanism should be differentially implemented in case of probabilistic visual stimulus presence and absence, and also resulted in distinct prediction error assessment. In our experimental protocol learned perceptual associations were assumed to occur exclusively because of repeated exposition to statistical regularity of sensory stimuli. To test our hypotheses, we examined neural activity during sensory conditioning by combining ERP analysis and multivariate classification. We then performed Pattern Similarity Analysis ^41,42^ to assess the relationship of activation pattern evoked by predictive auditory stimulus with that evoked by expected visual stimulus presentation or omission, by computing the cross-validated Mahalanobis distance (cvMD) ^43^. Finally, we modelled the evolution of prediction error signals during implicit learning of incidental perceptual regularities ^44,45^ using the Hierarchical Gaussian filter (HGF) modeling, a computational approach that was shown to successfully explain associative learning in terms of probabilistic perceptual priors ^12^. The HGF, a Bayesian ideal observer model for characterizing inferences of uncertain perceptual inputs ^46^, allowed us to estimate dynamic changes of prediction error signals in relation to differential probabilistic occurrences of visual input presentation and omission during sensory conditioning.

**Figure 1.**
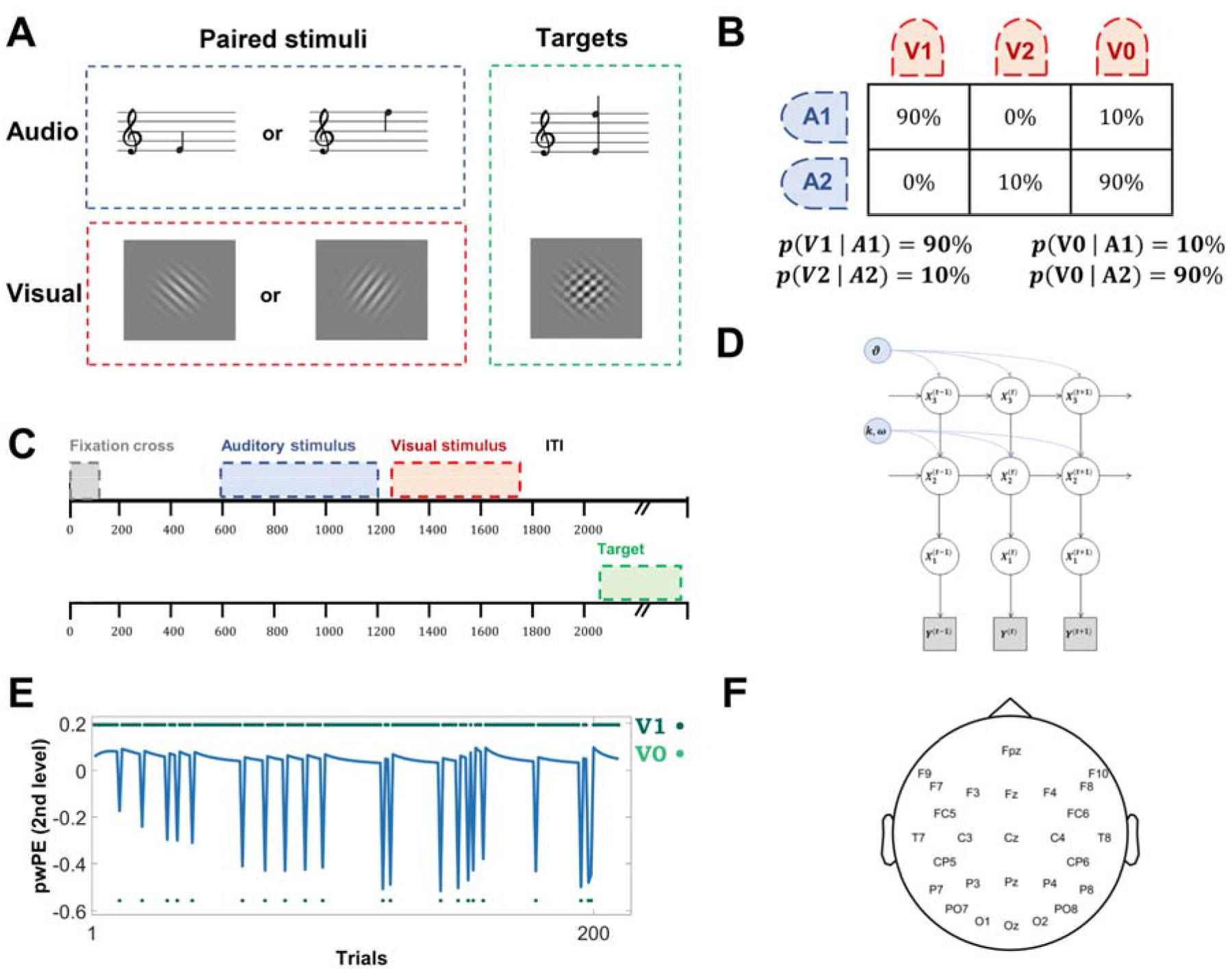
**A**. The experimental procedure entailed passive exposition to a stream of task-irrelevant auditory stimuli, A1 and A2 consisting of low and high frequency pure tones, 250 Hz and 500 Hz respectively (counterbalanced across participants), paired with task-irrelevant visual stimuli V1 and V2 consisting of two white Gabor patches with 45° and 135 orientation, presented against a grey background. The main task entailed detection and button press response only to two specific targets consisting of an auditory stimulus combining A1 and A2, and a visual stimulus combining V1 and V2. **B**. Contingency table showing the probabilistic occurrences of visual stimuli presentation given each auditory stimulus, and resulting conditional probabilities of the four different trials. **C**. Trial structure implied fixation cross presentation for 100 ms, followed after 500 ms by the presentation of one of two equally-probable auditory stimuli for 600ms, and after 50 ms of its offset by the presentation of one of two Gabor patches for 500 ms. Four target stimuli were also interleaved in a block after the ITI period. The representation of two trials’ time course shows that the target stimulus (lower line) was not always preceded by the audio-visual association (upper line). **D**. Graphical description of the Hierarchical Gaussian Filter (HGF) model adopted to depict individual trajectories of precision-weighted prediction error on the basis of ongoing variability in weighting between sensory evidence and perceptual beliefs. **E**. Exemplary single subject precision-weighted prediction error trajectory in relation to V1,V0|A1 condition. **F**. The EEG montage we adopted showing the position of the 27 channels.

## 2. RESULTS

We collected EEG and behavioral data from 21 human volunteers exposed to a stream of non-target auditory and visual stimuli and involved in a main audio-visual detection task implying button press responses to target stimuli presentation (Fig. 1A). Crucially, we manipulated the transition probabilities between the non-target stimuli so that non-target audio-visual co-occurrences had no predictable effects on target presentation, thus making the learning of these associations task-irrelevant (Fig. 1B-C). Participants debriefed at the end of the experiment reported not to be consciously aware of stimuli pairings. When specifically interrogated about possible stimuli associations they reported not to have noticed either regularities of stimuli presentation or specific audio-visual pairings.

### 2.1. Event-related potentials

We performed conventional event-related potential (ERP) analysis on both audio and post-audio periods considering the *initial* and *final* trials so as to assess the evoked activity throughout the learning of statistical associations of non-target auditory and visual stimuli. Results of ERP analysis comparing *initial* versus *final* trials of sensory conditioning in relation to audio period revealed a significant attenuation of signal amplitude in response to A1 auditory stimulus, predictive of V1 visual stimulus (V1|A1), in the interval 190 - 280 ms in the occipital ROI (Fig. 2A, *d* = 0.92, *BF*_10_ = 78.31, *p* = 0.0045, cluster corrected) and in response to A2 auditory stimulus, predictive of V0 stimulus (stimulus absence), in the interval 180 - 240 in the temporo-parietal ROI (Fig. 2A, *d* = 0.79, *BF*_10_ = 22.97, *p* = 0.0278, cluster corrected). Considering only *final* trials, a reduced negativity was observed for A1 with respect to A2 in the interval 180 - 230 ms in the frontal ROI (Fig. 2A, *d* = 0.77, *BF*_10_ = 19.57, *p* = 0.0260, cluster corrected) and in the interval 180 - 285 ms in the temporo-parietal ROI (Fig. 2A, *d* = 1.02, *BF*_10_ = 211.36, *p* = 0.008, cluster corrected). In the post-audio period, comparison of *initial* and *final* trials revealed a significant attenuation of the response to V1|A1 in the interval 60 - 170 ms in the frontal ROI (Fig. 2B, *d* = 0.84, *BF*_10_ = 38.03, *p* = 0.0155, cluster corrected) and in the intervals -10 - 160 ms (Fig. 2B, *d* = 1.11, *BF*_10_ = 443.03, *p* = 0.0134, cluster corrected) and 530 - 600 ms (Fig. 2B, *d* = 0.64, *BF*_10_ = 5.82, *p* = 0.0236, cluster corrected) in the parieto-occipital ROI. A significant signal attenuation was also observed for V0|A2 condition in the interval -50 - 40 ms in the frontal ROI (Fig. 2B, *d* = 0.69, *BF*_10_ = 9.25, *p* = 0.0466, cluster corrected) and in the interval -30 - 30 ms in the parieto-occipital ROI (Fig. 2B, *d* = 0.75, *BF*_10_ = 15.95, *p* = 0.0321, cluster corrected). In short, these results revealed across conditioning changes of neural responses to both predictive auditory and predicted visual stimuli, that is the acquired conditioned response and unconditioned response, respectively.

**Figure 2.**
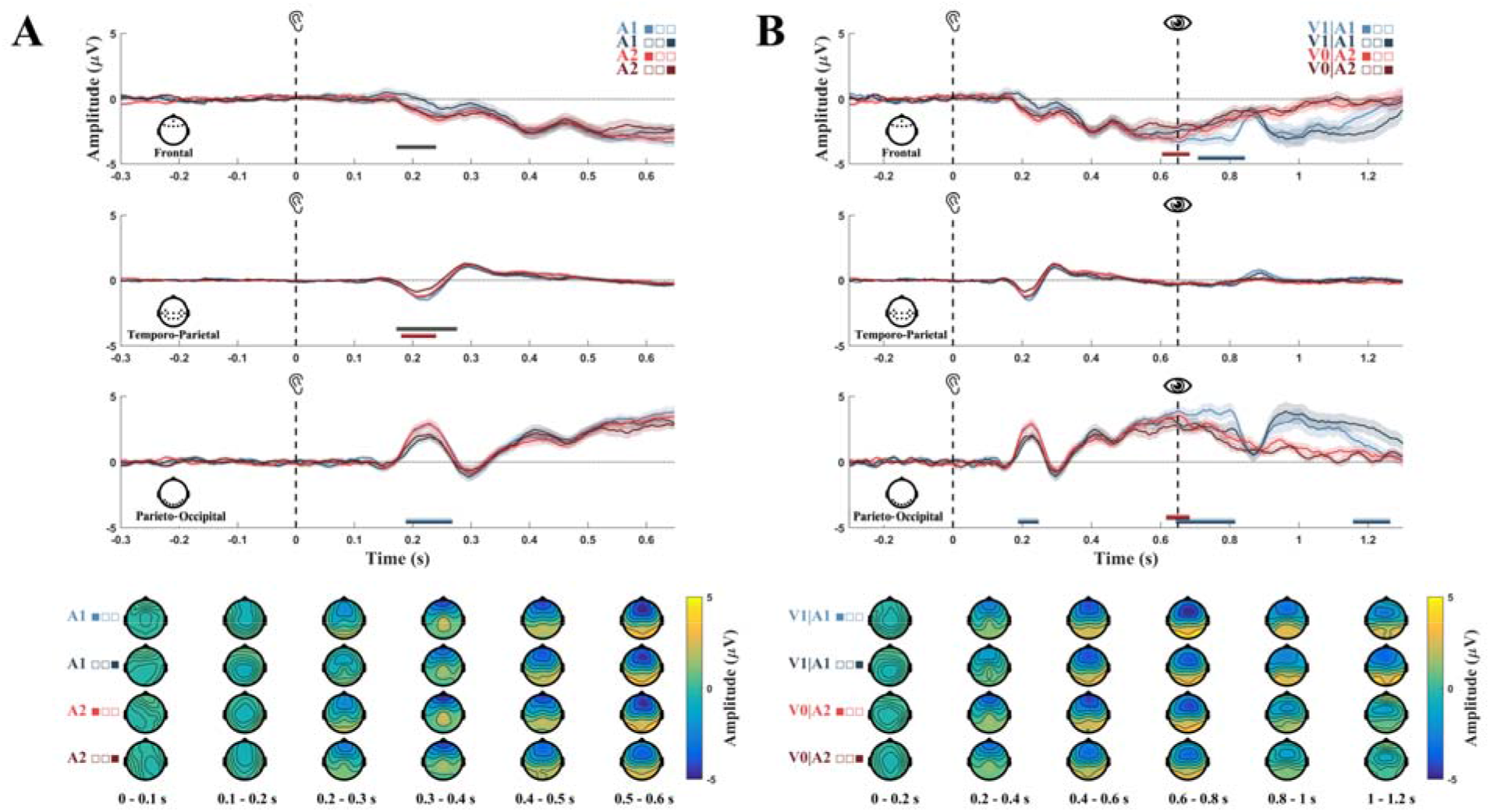
**A**. Top. ERPs of initial (A1 light blue, A2 light red) and final (A1 dark blue, A2 dark red) trials in relation to auditory stimuli onset for each selected ROI. Shading refers to SEM across participants, horizontal bars refer to statistical significance.Bottom, topographic maps depicting whole-brain spatial distribution of EEG signal across 100ms intervals after auditory stimuli onset. **B**. Top. Full time-course ERPs of initial (V1|A1 light blue, V0|A2 light red) and final (V1|A1 dark blue, V0|A2 dark red) trials including both auditory and visual stimuli presentation for each selected ROI. Bottom. Topographic maps depicting whole-brain spatial distribution of EEG signal across 100ms intervals after auditory stimuli onset.

### 2.2. Multivariate classification

Multivariate decoding, performed using Linear Discriminant Analysis-based classification, aimed to assess EEG signal differences between A1 and A2 during the whole conditioning session. Such expected differences should reflect differential processing of equivalent auditory stimuli that distinctively anticipated visual stimulus presence or absence. Our results showed significant discrimination performance between A1 and A2 with respect to chance level in three different time windows (Fig. 3A): 192-340 ms (*AUC*_*peak*_ = 0.56, *d* = 1.17, *BF*_10_ = 771.98, *p* < 0.0001, cluster corrected), 344-444 ms (*AUC*_*peak*_ = 0.53, *d* = 0.85, *BF*_10_ = 39.42, *p* = 0.0017, cluster corrected) and 500-540 ms (*AUC*_*peak*_ = 0.52, *d* = 0.73, *BF*_10_ = 13.04, *p* = 0.0484, cluster corrected), that is immediately preceding probabilistic visual stimulus presence or absence. These results thus indicated distinct processing of equivalent A1 and A2 stimuli differentially predicting V1 and V0 respectively. Relevant channels for classification were located in the frontal regions for the first significant time window and mostly in the temporo-occipital regions for the other two time windows.

**Figure 3.**
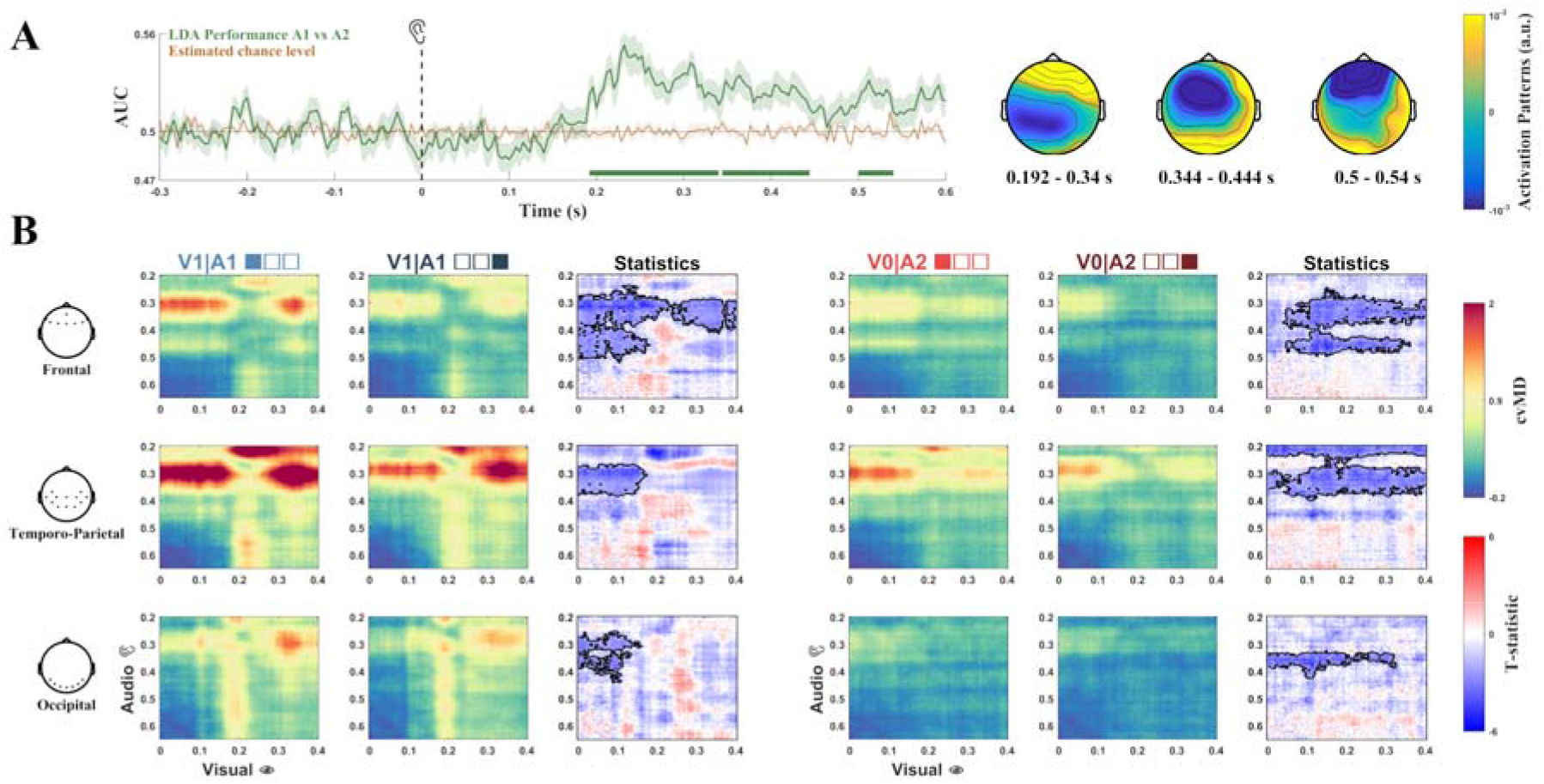
**A**. Left, EEG signals classification performance (Area Under the Curve) of A1 versus A2 during audio period (green) against estimated chance-level (orange). Shading indicates SEM across folds. Green horizontal lines indicate statistical significance. Right, topographic maps of activation patterns computed from the model’s parameters, representing feature importance for classification performance in relation to EEG channels and time windows. **B**. Pattern Similarity Analysis. PSA aimed to test whether the pattern elicited by the predictive auditory stimulus increasingly resembled the pattern elicited by the expected visual stimulus during sensory conditioning. The images show Pattern Dissimilarity Matrices of EEG signal considering audio and post-audio periods for V1|A1 (left) and V0|A2 (right) condition, for each selected ROI. For each condition, the first column refers to cvMD during initial trials, the second column refers to cvMD during final trials, and the third column depicts statistical differences between initial and final trials. Outlined regions indicate statistical significance. X and Y axes depict time course of visual and auditory stimulus respectively. For the visual stimulus the temporal interval ranges between visual stimulus onset and its offset, whereas for the auditory stimulus it ranges between 200 ms after auditory stimulus onset (this time was set on the basis of ERP and MVPA classification results) and its offset.

### 2.3. Pattern similarity analysis

Pattern similarity analysis aimed to test whether the pattern elicited by the predictive auditory stimulus increasingly resembled the pattern elicited by the expected visual stimulus during sensory conditioning. Pattern Similarity Analysis (PSA) (was performed by calculating a dissimilarity matrix (DM) between audio and post-audio periods to specifically assess the relationship of brain responses evoked by predictive A1 and A2 stimuli with those evoked by predicted V1 and V0, respectively (Fig. 3B). DM, estimated on the basis of the cross-validated Mahalanobis distance (cvMD), showed that in the V1|A1 condition the cvMD was significantly lower in the *final* trials with respect to *initial* trials in several clusters in frontal, temporo-parietal and occipital ROIs. In the frontal ROI two main significant clusters were observed, one corresponding to ∼300-500 ms in the audio period and ∼0-200 ms in the post-audio period (*d* = 1.36, *BF*_10_ = 4608.53, *p* < 0.0001, cluster corrected), and another one corresponding to ∼300-400 ms in the audio period and ∼250-400ms in the post-audio period (*d* = 0.84, *BF*_10_ = 36.18, *p* = 0.0009, cluster corrected). In the temporo-parietal (*d* = 1.02, *BF*_10_ = 202.98, *p* = 0.005, cluster corrected) and parieto-occipital (*d* = 0.99, *BF*_10_ = 161.58, *p* = 0.015, cluster corrected) ROIs the significant cluster corresponded to ∼300-400 ms in the audio period and ∼0-150 ms in the post-audio period. In V0|A2 condition, the cvMD was also significantly lower in the *final* trials with respect to *initial* trials in several clusters in frontal, temporo-parietal and occipital ROIs. In the frontal ROI two main significant clusters were observed, one corresponding to ∼300-400 ms in the audio period and ∼50-400 ms in the post-audio period (*d* = 1.07, *BF*_10_ = 313.13, *p* = 0.0048, cluster corrected), and another one corresponding to ∼400-500 ms in the audio period and ∼50-350ms in the post-audio period (*d* = 1.21, *BF*_10_ = 1132.09, *p* = 0.005, cluster corrected). In the temporo-parietal ROI two significant clusters were also observed, one corresponding to ∼200-250 ms in the audio period and ∼0-400 ms in the post-audio period (*d* = 1.10, *BF*_10_ = 434.36, *p* = 0.002, cluster corrected), and another one corresponding to ∼280-380 ms in the audio period and ∼0-500ms in the post-audio period (*d* = 1.03, *BF*_10_ = 219.11, *p* = 0.0019, cluster corrected). In addition, a significant cluster was observed in the occipital ROI corresponding to ∼320-400 ms in the audio period and ∼0-300ms in the post-audio period (*d* = 1.21, *BF*_10_ = 1124.35, *p* = 0.007, cluster corrected). Altogether these results showed decreased dissimilarity for both V1|A1 an V0|A2 conditions during sensory conditioning.

### 2.4. HGF modelling

We modelled individual trajectories of precision weighted prediction error (pwPE) on the basis of ongoing variability in weighting between sensory evidence and perceptual beliefs using the Hierarchical Gaussian filter (HGF), a Bayesian ideal observer model that attempts to predict future stimuli occurrence on the basis of the history and uncertainty of contextual events. This analysis allowed us to assess differential prediction error processing associated with violation of expected presentation or omission of visual stimulus, and to further demonstrate that associative learning of perceptual stimuli can occur even when perceptual stimuli are task-irrelevant. We thus fed into the HGF model the same sequence of stimuli that every subject was exposed to, and then we obtained the prediction error trajectory. We used these ideal error trajectories as regressor in a general linear model (GLM) applied to each channel-time point pair, for each participant. HGF analysis showed that pwPE, significant for both V1,V0|A1 and V2,V0|A2, was mediated by all selected brain regions, although the largest effect size was in the occipital ROI (Fig. 4). Regression analysis resulted in a significant effect for both V1,V0|A1 and V2,V0|A2 (Fig. 4) in the interval ∼250-550 ms of post-audio period in the frontal (V1,V0|A1: *d* = 0.66, *BF*_10_ = 7.29, *p* = 0.0007, cluster corrected; V2,V0|A2: *d* = 0.72, *BF*_10_ = 11.84, *p* = 0.0007, cluster corrected) and temporo-parietal ROIs (V1,V0|A1: *d* = 0.81, *BF*_10_ = 26.01, *p* = 0.0003, cluster corrected; V2,V0|A2: *d* = 0.69, *BF*_10_ = 8.89, *p* = 0.0005, cluster corrected), and ∼150-550 ms in the occipital ROI (V1,V0|A1: *d* = 0.72, *BF*_10_ = 11.96, *p* = 0.0386, cluster corrected; V2,V0|A2: *d* = 0.77, *BF*_10_ = 18.89, *p* < 0.0001, cluster corrected). Finally, ***R***^2^was significantly larger for V1,V0|A1 than for V2,V0|A2 (Fig. 4) in the interval ∼370-400 ms in the post-audio period in the occipital ROI (*d* = 0.64, *BF*_10_ = 6.03, *p* = 0.0398, cluster corrected), indicating differential neural processing of PE in relation to violation of predicted visual stimulus presentation and omission.

**Figure 4.**
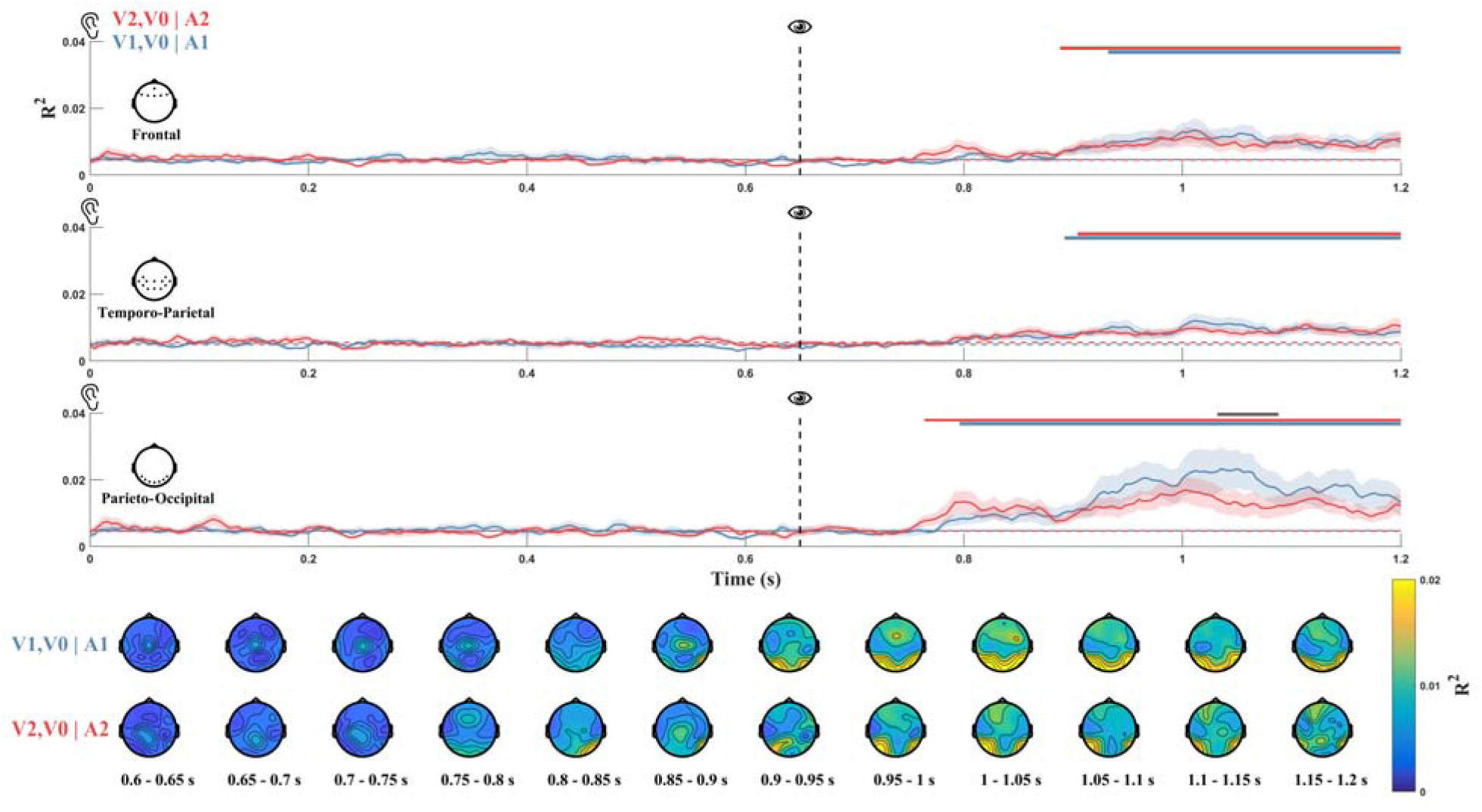
Top. Line plots of GLM performance (*R*^2^) fitting HGF model-derived precision-weighted prediction error trajectories to each EEG channel, averaged across ROIs, for V1,V0|A1 (solid blue) and V2,V0|A2 (solid red) conditions. Dotted lines represent average performances during baseline period (preceding the onset of visual stimuli). Shading indicates SEM across participants. Horizontal lines indicate statistical significance of differences between each model and baseline (blue and red for V1,V0|A1 and V2,V0|A2 respectively), and between V1,V0|A1 and V2,V0|A2 (grey). Bottom. Topographic maps depicting whole-brain spatial distribution of GLM performance for each model across 50ms intervals in the post-audio period.

## 3. DISCUSSION

In our study, we investigated whether incidental but recurrent exposition to non-salient and task-irrelevant probabilistic audio-visual associations induced implicit learning of perceptual patterns resulting in conditioned cortical responses representing early instantiations of predicted stimulus-related sensory information. To this aim, we assessed both temporal and predictive aspects of neural signals associated with learned conditional probability of paired audio-visual stimuli. Our analyses showed that incidental perceptual regularities were rapidly encoded leading to differential neural responses to anticipatory auditory stimuli predictive of either visual stimulus presence or absence. ERPs results revealed that in the final trials of sensory conditioning brain responses to equivalent A1 and A2 stimuli, anticipating probabilistic visual stimulus presence and absence respectively, were significantly different about 200 ms after stimulus onset, in both frontal and temporo-parietal regions (Fig. 2A-B). Specifically, a response attenuation in the temporo-parietal region was observed in response to A2 but not to A1, suggesting that A1, anticipating V1 presentation in contrast to A2 preceding visual stimulus absence, required differential processing. In line with the known audiovisual cross-modal effect resulting in visual cortex activity during auditory perception ^47,48^, increased signal amplitude in the parieto-occipital region was observed in response to both A1 and A2 across all trials. Though, a significant attenuation over time in the parieto-occipital region was measured in response to A1, but not to A2. The repetition suppression effect observed only for A1 points to differential predictive processes in relation to expected visual stimulus presentation and omission.

According to the assumption that expectation suppression effect reflects decreased surprise signal we would have expected to observe a similar attenuation for both A1 and A2. Vice versa, in line with the interpretation of sensory attenuation resultant of sharpening of stimulus representation ^2^, we interpreted the parieto-occipital differential response to A1 with respect to A2 as increased response tuning, as A1 in contrast to A2 carries information about consecutive V1 presentation. Our result are also in line with previous evidence of enhanced stimulus-specific baseline activity during early sensory processing in relation to probabilistic signal occurrence ^13^. Multivariate pattern analysis corroborated ERP results showing differential neural responses to A1 and A2 in particular in a temporal interval immediately preceding probabilistic visual stimulus presence or absence, and disclosed that such divergent responses manifested early in the frontal region and later in the temporo-occipital areas (Fig. 3A). These results might also potentially ascribable to the inherent difference of auditory stimuli, though the feature importance, showing that occipital channels were particularly relevant in later stages of the classification (350-600 ms), suggests that the two auditory stimuli might indeed convey differential predictive information of consequent stimulus presentation. Pattern similarity analysis revealed that for V1|A1 condition the cvMD between audio and post-audio period, initially large, decreased over time in the frontal, parieto-occipital and temporo-parietal regions, resulting in decreased dissimilarity of neural activity between ∼0-150ms after V1 onset and ∼250-400ms after A1 onset (Fig. 3B). Decreased cvMD was also observed in the frontal region between neural activity ∼400-500ms after A1 onset and ∼0-150ms after V1 onset (Fig. 3B).

Altogether, our findings revealed that over time the pattern of neural activity elicited by A1 increasingly resembled that elicited by V1, and thus indicated that some specific perceptual priors relative to probabilistic but task-irrelevant audio-visual association were instantiated at cortical level during the unfolding of sensory conditioning. Previous studies showed that prior expectations can elicit anticipatory sensory-like neural representations of predictable sensory information. In particular, Kok and colleagues reported that a predictive auditory cue evoked early sensory representation in the primary visual cortex immediately before the expected visual stimulus ^28^. In our study, in line with previous findings showing expectation-induced decrease of primary visual cortex activity ^8,9,24,49^, we also measured an attenuated response over time during V1|A1, and immediately preceding V1, in the parieto-occipital and frontal regions, but similarity analysis did not show evidence of pre-stimulus neural instantiation of expected sensory input.

Notably, the cvMD decreased over time also in relation to V0|A2 condition, in the frontal, parieto-occipital and temporo-parietal regions. In particular, decreased A2-related cvMD in the temporo-parietal region was observed between neural activity around 200-250ms and 280-350ms in the audio period, and that in the V0-V2 post-audio period (0-400ms). These results are possibly ascribable to the attenuated response to A2, resulting in increased similarity with the response to frequent V2 omission. On the other hand, as for A1 stimulus, these effects might also reflect tuning of A2 response progressively incorporating perceptual priors of V2 occurrence’s low probability.

Accordingly, decreased dissimilarity was also observed in the parieto-occipital region between neural activity occurring 320-400 ms after A2 and during V0-V2 time period. Moreover, neural attenuation over time was also observed for V0|A2 condition, in both frontal and parieto-occipital regions, in a time window anticipating and also corresponding to V2 presentation, further suggesting increased encoding of low V2 occurrence probability.

Overall, our results indicated that forward neural processes anticipating some specific aspects of consecutive visual stimulus presentation might occur earlier during initial processing of predictive stimulus; however, due to limited spatial resolution the exact representational content of such anticipatory activity remains to be fully clarified.

As reported in previous studies ^9,19^ we observed anticipatory neural mechanisms independently of stimulus salience, as our stimuli consisted of non-salient abstract and auditory and visual stimuli. In addition, the observed effects were task-independent, since statistical regularity of paired audiovisual stimuli was not necessary and irrelevant to detect target stimuli. This result is also in line with studies showing that task-irrelevant visual perceptual learning can occur as a result of mere exposure to perceptual features.

It has been proposed that several aspects of perceptual and statistical learning might be unified in the framework of Hierarchical Bayesian framework ^50^. In both perceptual and statistical learning attention plays an important role ^34,51^. In visual perceptual learning attention enhances bottom-up signals from task-relevant features, whereas it decreases signals from task-irrelevant features; however, visual perceptual learning of task-irrelevant features can also occur as far as they can be optimally attended (suprathreshold presentation) ^51^. Similarly, in statistical learning attenuation of neural response to predictable stimuli vanishes when they are not attended ^34^. In our study, as non-target stimuli shared both auditory and visual features with target stimuli, we supposed that they were likely attended, however overt encoding of the statistical regularity of audio-visual associations was not necessary being completely irrelevant to the task.

The observed conditioned responses in the sensory cortices appeared to be mediated by frontal and prefrontal areas. The differential responses to A1 and A2 in the frontal ROI, preceding changes in temporo-occipital ROIs, suggest that prefrontal regions might support mutual information exchange between auditory and visual cortices ^52^, likely in relation to temporal aspects of perceptual regularities ^53,54^, for timely instantiation of specific perceptual priors. Sensory priors inducing auditory-cued shaping of visual cortex responses in both V1|A1 and V0|A2 conditions might be then mediated by direct interactions between auditory and visual cortices ^49^. Remarkably, computational modeling of pwPE trajectories further demonstrated instantiation of predictive neural signals by showing distinct neural processing of prediction error in relation to violation of expected visual stimulus presence or absence. Moreover, differential neural processing of PE correlated with activity in frontal, temporal, and occipital areas (Fig. 4) at latencies corresponding to those of typical event-related potentials elicited by deviant stimuli ^55,56^. Prediction’s precision of audiovisual patterns, likely mediated by temporal regions such as the superior temporal gyrus ^57,58^, might trigger gradual update of prefrontal regions-mediated cortical representations of expected V1 and V0 ^59,60^. Indeed, analysis of pwPE trajectories revealed a significant difference of V1|A1 with respect to V0|A2 occurring about 300-400 ms after V1 and V0 onset in the occipital region, indicating stimulus-specific differential processing of prediction violation ^61,62^.

## 4. CONCLUSION

Our study demonstrated rapid encoding of incidental but probabilistic audio-visual regularities leading to modulation of cortical responses to both predictive and predicted sensory stimuli. Sensory conditioning of task-irrelevant audio-visual associations also appeared to induce increased similarity between the neural response to predictive auditory stimuli and the response to predicted visual stimulus presence or absence. Remarkably, Hierarchical Gaussian filter modeling estimating dynamic changes of prediction error signals in relation to differential probabilistic occurrences of audio-visual stimuli further demonstrated instantiation of predictive neural signals by showing distinct neural processing of prediction error in relation to violation of expected visual stimulus presence or absence. Overall, our findings suggest that statistical learning of non-salient and task-irrelevant perceptual regularities might induce generation of neural priors at the time of predictive stimulus presentation conveying sensory-specific information of the predicted consecutive stimulus. As in the case of goal-directed behavior where predictive mechanisms were related to information-seeking random exploration ^60,63^, the intrinsic neural need of reducing uncertainty about state transition dynamics of the environment might also explain learned incidental probabilistic sensory patterns. Learning sensory representations might then depend on automatic perceptual processes exploiting either reward statistics of past experience or beliefs about future representations ^64^ that optimize neural computations for adaptive behavior.

Ultimately, in accordance with predictive brain principles our results suggest that associative processes might even occur exclusively at perceptual level, possibly as a consequence of Hebbian neural plasticity ^65^, and that stimulus salience that is typically considered a critical element for learning in classical conditioning might be in fact not strictly required ^66,67^.

## 5. METHODS

### 5.1. Participants

Twenty-one volunteers (13 females, range 19-32, mean age 24.3 ± 3.4 (SD)) participated in the study. All were right-handed with normal or corrected-to-normal vision and normal hearing, had no history of neurological disorders and were not taking any neurological medications. All participants gave written informed consent. The study was conducted in accordance with the Declaration of Helsinki and approved by the Ethics Committee of the University of Trento.

### 5.2. Procedure

During the experimental procedure participants were exposed to a stream of auditory and visual stimuli while sitting in a dimly-lit booth at a distance of 1 m from the monitor (22.5” VIEWPixx; resolution: 1024 × 768 pixels; refresh rate: 100 Hz; screen width: 50 cm). Participants were informed to be involved in an audio-visual detection task implying button response only to target stimuli presentation. Auditory stimuli A1 and A2 consisted of low and high frequency pure tones, 250 Hz and 500 Hz respectively, whereas visual stimuli V1 and V2 consisted of two white-colored Gabor patches with 45° and 135 orientation (4.4° × 3.4° visual angle, generated with Gaussian envelope, standard deviation = 18.0, spatial frequency = 0.08 cycles/pixel), presented against a grey background (Figure 1A). In each trial auditory stimuli were followed by the presentation of visual stimuli according to an equivalent temporal sequence with two opposite probability distributions resulting in high frequent and low frequent visual stimulus occurrence (Figure 1B). In the condition of high frequent visual stimulus occurrence, A1 stimulus was followed by V1 Gabor patch (V1|A1) 90% of the time and 10% by visual stimulus absence, V0|A1. In the condition of low frequent visual stimulus occurrence, A2 stimulus was followed by a visual stimulus (V2|A2) 10% of the time and by visual stimulus absence 90% of the time, V0|A2. Pairings of the auditory and visual stimuli were counterbalanced across participants. Each trial started with a fixation cross presented for 100 ms, followed after 500 ms by the presentation of one of two equally-probable auditory stimuli for 600ms, and after 50 ms of its offset by the presentation of one of two Gabor patches for 500 ms (Figure 1C). Trials of V0|A1 and V0|A2 conditions, which were of equal length, entailed no Gabor patch presentation. Trials were interspersed with an inter-trial interval (ITI) of 2500 ms ± 500 ms. The main audio-visual target detection task implied button press response only at the presentation of specific target stimuli represented by an auditory target combining A1 and A2 stimuli, and a visual target combining V1 and V2 stimuli, both lasting 500ms and followed by equal ITI. The experimental session consisted of 400 trials presented in 10 blocks, each including random presentation of 4 perceptual targets, with a total duration of about 40 min. The experimental protocol was implemented using OpenSesame and PsychoPy as backend ^68^. The experimental procedure aimed at inducing more attentional resources allocation to task-relevant perceptual targets. Importantly, probabilistic contingencies of audio-visual pairing were completely irrelevant to the audio-visual target detection task.

### 5.3. EEG data acquisition and preprocessing

EEG data were recorded with a standard 10-5 system and 27 Ag/AgCl electrodes cap (EasyCap, Brain Products, Germany) at a sampling rate of 1 kHz. Impedance was kept below 10 kΩ for all channels. AFz was used as the ground and the right mastoid was used as reference. Electrodes were approximately evenly spaced and positioned at the following scalp sites: Fpz, Fz, F3, F4, F7, F8, F9, F10, FC5, FC6, T7, C3, Cz, C4, T8, CP5, CP6, P7, P3, Pz, P4, P8, PO7, PO8, O1, Oz, and O2 (Fig. 1F). All preprocessing steps were conducted using EEGLAB ^69^ in accordance with guidelines and recommendations for EEG data preprocessing such as HAPPE ^70^, that are also applicable in the case of low-density recordings, HAPPILEE ^71^. Spherical interpolation was carried out on a limited number of bad channels on the basis of channel correlation lower than 0.85 on average with respect to its neighbours and guided by visual inspection (average number of interpolated channels: 0.74, range: 0-3). Data were down-sampled at 250 Hz, high-pass filtered at 0.1 Hz and low-pass filtered at 80 Hz, using a Butterworth IIR filter with model order 2. CleanLine (https://github.com/sccn/cleanline) with default parameters was used to remove power line 50Hz noise and its harmonics up to 200 Hz. Data were then re-referenced to a common average reference ^71^ and epoched between -300 ms and 1300 ms relative to the onset of the auditory stimulus with a baseline correction between -300 ms and 0 ms. Artifact rejection was performed through visual inspection and by an automatic procedure excluding epochs with very large signal amplitudes (detection threshold = ±500). The average number of trials rejected per participant was 1.1% (SD=2.1%, range 0-7.3%). Stereotypical artefacts, including eyeblinks, eye movements and muscle artefacts, were detected via independent component analysis using the extended Infomax algorithm ^72^. A rejection strategy based on ICLabel ^73^ and visual inspection resulted in the removal of an average number of independent components equal to 9.33 (±3.48 SD). Finally, data were converted to Fieldtrip format ^74^ for subsequent analyses.

### 5.4. EEG data analysis

Data analysis aimed at assessing implicit associative learning of paired audio-visual stimuli by investigating neural responses evoked by predictive auditory stimuli and by predicted visual stimulus presentation or omission. The analysis focused on two main epochs by performing grand averaging considering separately auditory and visual stimuli: 0 - 650 ms relative to auditory stimuli presentation (audio period), and 650 - 1300 ms relative to Gabor patch presence or absence (post-audio period). Trials were divided into three equivalent groups to analyze *initial* (first third of trials) and *final* phases (last third of trials) of sensory conditioning, and each phase consisted of 54 trials (except possible rejection of trials due to artifacts). Conventional event-related potential (ERP) analysis was first performed on both audio and post-audio periods considering *initial* and *final* trials. All trials of A1, A2, V1|A1 and V0|A2 conditions were averaged separately in relation to *initial* and *final* phases, whereas trials of V0|A1 and V2|A2 conditions were not analyzed due to their limited number required by the specific contingencies schedule of our conditioning paradigm. As in previous related studies ^75,76^, for ERP analysis we adopted an approach that considers the average neural response over comparable predefined regions of interest (ROI) using frontal (Fpz, Fz, F3, F4, F7, F8, F9, F10), temporo-parietal (FC5, FC6, T7, C3, Cz, C4, T8, CP5, CP6, P7, P3, Pz), and parieto-occipital channels (P4, P8, PO7, PO8, O1, Oz, O2), and thus enables more robust statistics through cluster-based correction. In addition, multivariate pattern classification based on Linear Discriminant Analysis (LDA) examining EEG signal differences between A1 and A2 at subject level was performed considering all conditioning trials and channels as samples and features respectively, with the MVPA-Light toolbox ^77^ and custom MATLAB scripts. Z-scoring was applied across samples for each time point separately to normalize channel variances and remove baseline shifts. A 5-fold cross validation scheme was adopted and the Area Under the Curve (AUC) was used as performance measure of LDA. An empirical chance level was obtained by running twice the same classification analysis with the same hyperparameter but with permuted labels. Statistical significance of model performance with respect to empirical chance level was assessed at group level (paired permutation t-test two tailed, α = 0.05) using mass univariate cluster-based permutation tests (10000 iterations) and maxsum as cluster statistic, a valid and powerful way of dealing with the problem of multiple comparisons ^78,79^. Effect size was estimated using Cohen’s d (*d*) and Bayes Factor t-test (*BF*_10_), reporting the peak value inside the significant cluster. Analysis of relevant channels for classification was performed converting the estimated weights of the LDA model at each fold into interpretable activation patterns ^80^. Furthermore, we performed pattern similarity analysis to estimate the similarity of brain responses evoked by predictive A1 and A2 stimuli with those evoked by expected V1 and V0, by calculating Pattern Dissimilarity Matrices based on the cross-validated Mahalanobis distance for each participant. A 5-fold cross validation scheme ^43^ was applied to assess the similarity of EEG signal between audio (200 - 650 ms) and post-audio (0 - 550 ms) periods, considering time points (EEG samples) of *initial* and *final* trials. These time windows were selected as a result of the ERP analysis and MVPA. Statistical significance of Dissimilarity Matrices (DM) differences between *initial* and *final* trials was tested at group level with cluster-based permutation tests with the same hyperparameters as described above.

### 5.5. Hierarchical Gaussian Filter modelling

The HGF is a Bayesian generative model ^46,81^ of perceptual inference on a changing environment based on sequential input ^12,82^. The HGF consists of perceptual and response models, representing a Bayesian ideal observer who receives a sequence of inputs and generates behavioral responses. Since our experimental design deliberately precluded behavioral responses, we used only the perceptual model ^83^. In this framework, a perceptual model comprised 3 hierarchal hidden states (*x*), which accounted for a multi-level belief updating process of the hierarchically related environmental states giving rise to sensory inputs, and the observed input (*y*) representing the actual occurrence of a stimulus in a given trial (Fig. 1D).

Our HGF model assumed that environmental hidden states evolved conditionally on the states at the immediately higher level. The hidden states processed at the first level of the perceptual model represented a sequence of beliefs 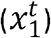 about stimulus occurrence, that is, whether a visual stimulus was presented (*y*^*t*^ = 1) or absent (*y*^*t*^ = 0) at trial *t*, and was modelled as follows:

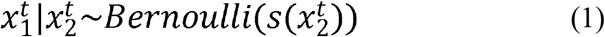

where 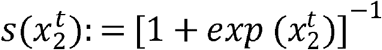 is the logistic sigmoid function. Here, the hidden states at the second level 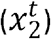 is an unbounded real parameter of the probability that 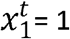, thus representing the current belief of the probability that a given stimulus occurred. Such a hidden state process evolves according to a Gaussian random walk:

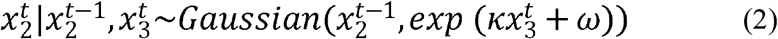

which depends on both its value at a previous trial *t*, and the hidden state at the third level of the hierarchy. In particular, the higher-level hidden state process 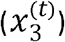 determines the log-volatility of the hidden state process at the second level, thus codifying the volatility of the environment during the time course of the experiment. This process evolves according to a Gaussian random walk:

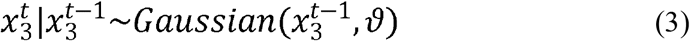

The parameter set (*k, ω, ϑ*) determined the dispersion of the random walks at different levels of the hierarchy and allowed us to shape individual difference in learning. By inverting the generative model, given a sequence of observations (*y*), it was possible to obtain the updating process of the trial-by-trial estimates of the hidden state variables.

The update rules shared a common structure across the model’s hierarchy: at any level *i* the update of the posterior mean 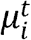 of the state *x*_*i*_, that represented the belief on trial *k*, was proportional to the precision-weighted prediction error (pwPE) 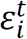 as follows:

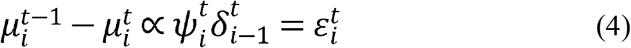

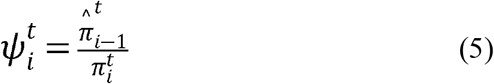

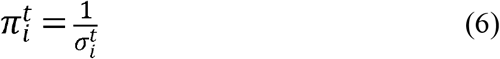

As shown in Eqs 4–6, in each trial, a belief update 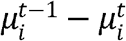 is proportional to the prediction error at the level below 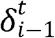. The pwPE is the product of the prediction error 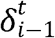 and a precision ratio 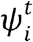 that depends on the precision (inverse variance, Eq. 5) of the prediction at the level below 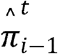 and the current level 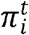. In this application, we were interested in the update equations of the hidden states at the second level, which have a general form similar to those of traditional reinforcement learning models, such as the Rescorla-Wagner model ^84^. The pwPE on the second level, was thus assumed to be responsible for the learned perceptual associations. The nature of the pwPE could be described through the following update equation of the mean of the second level:

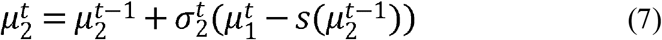

where the last term represents the prediction error 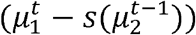 at the first level weighted by the precision term 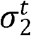 (see ^46^ for a general derivation and more mathematical details). Individual trajectories of pwPEs with separate models for V1,V0|A1 and V2,V0|A2 were calculated by estimating the parameters that minimized Bayesian Surprise using the Broyden-Fletcher-Goldfarb-Shannon (BFGS) quasi-Newton optimization algorithm. We determined these Bayes optimal perceptual parameters by inverting the perceptual model based on the stimulus sequence alone and a predefined prior for each parameter (HGF toolbox, version 5.2 implemented via the Translational Algorithms for Psychiatry Advancing Science toolbox). These model-derived trajectories of pwPEs (Fig. 1E) from the second level were used as regressor in a general linear model (GLM) applied to each channel-time point pair for each participant. We used the *R*^2^ measure for evaluating the goodness of fit and averaged these values over the selected ROIs. Statistical significance was tested at group level with cluster-based permutation tests and using the same hyperparameters as previously described.

## AUTHOR CONTRIBUTIONS

AG: Conceptualization, Methodology, Software, Formal analysis, Investigation, Data curation, Visualization, Writing-original draft, Writing - Review & Editing

MD: Software, Formal analysis, Writing - Review & Editing

GG: Investigation, Writing - Review & Editing

CR: Software, Formal analysis, Writing - Review & Editing

CB: Methodology, Writing-Review & Editing

AC: Conceptualization, Methodology, Writing-original draft, Writing - Review & Editing, Supervision, Project administration

## ETHICS STATEMENT

The study was approved by the ethics committee at the University of Trento (protocol n° 2018-009) and conformed to the Declaration of Helsinki.

## ACKNOWLEDGMENTS

None.

